# A transcriptome atlas of *Striga hermonthica* germination

**DOI:** 10.1101/2022.12.14.520245

**Authors:** Gilles Irafaha, Sylvia Mutinda, Fredrick Mobegi, Brett Hale, George Omwenga, Asela J. Wijeratne, Susann Wicke, Emily S. Bellis, Steven Runo

## Abstract

**Societal Impact Statement:** Witchweeds, parasitic plants of the genus *Striga*, are nicknamed “cereal killers” because of their devasting destruction of Africa’s most staple cereals, including maize, sorghum, millets, and upland rice. The parasite relies on biomolecules emitted from the host roots to germinate and therefore initiate its infectious lifecycle. Some sorghum varieties have evolved to not produce effective germination stimulants, making them resistant to the parasite. Here, we assess genetic factors that underpin *Striga* germination. We discuss how such knowledge can be used to develop new *Striga* management strategies through the disruption of host-parasite communication exchange.

**Summary:**

- Seeds of the parasitic plant *Striga* are dormant. They only germinate in response to biomolecules emitted from the host’s root exudate, strigolactones (SL). But, it is now emerging that *Striga* germination is a much more complex process regulated by crosstalk of hormone signaling pathways.
- To further understand the genetic basis of the communication exchange between *Striga* and its host sorghum, we performed a comparative transcriptomic analysis. We sought to identify major transcriptomic changes that define the germination process in *Striga* and a set of genes that may contribute to the differences in germination rates.
- Results showed that germination proceeds immediately after SL perception and is marked by a wave of transcriptional reprogramming to allow for metabolic processes of energy mobilization. Cluster analysis using self-organizing maps (SOMs) revealed a time-phased and genotype-differentiated response to germination stimulation. The variation in germination was also a function of hormonal crosstalk. The early germination stage was associated with significant repression of genes in the abscisic acid (ABA) biosynthesis pathway. Other hormones influenced germination as follows: (i) ABA and auxin repressed germination, (ii) brassinosteroid, ethylene and jasmonic acid promoted germination, and (iii) cytokinin had a more prominent role post-germination rather than during germination. Perception of SL sets the germination programme leading to different rates of germination in sorghum followed by a complex hormonal regulation network that acts to either repress or enhance germination. These results have far-reaching implications for developing *Striga* management strategies by disrupting hormonal communication exchange.

## 1. INTRODUCTION

*Striga* is a genus of root parasitic plants belonging to the family Orobanchaceae. Parasitism occurs when the parasite seed germinates and attaches to the roots of the host to extract water and photoassimilates. Soon after attachment, the host wilts, and suffers irreversible growth retardation or death (Hearne, 2009). The parasite is most prevalent in Sub–Saharan Africa (SSA), where two of its most destructive species *S. asiatica* (L.) Kuntze and *S. hermonthica* (D.) Benth greatly limit production of the most staple cereals including maize, millets, sorghum, and upland rice, exposing millions to hunger and starvation. *Striga* infests approximately 40 million hectares of arable land and result in 30 to 100 percent losses (reviewed in: Spallek et al., 2013). Most control strategies have achieved low to moderate success as the parasite continues to thrive and expand its host and natural range, becoming one of the most intractable problems of African agriculture (Babiker, 2007).

*Striga*’s highly successful parasitic lifestyle can be largely attributed to its remarkably adapted seeds (Runo and Kuria, 2018). Each *Striga* plant produces hundreds of thousands of tiny dusty seeds that are deposited in the soil every planting season (Berner, 1995). The seeds stay dormant in the soil and germinate after conditioning under warm temperatures and moisture, a cue for suitable environmental conditions. Following conditioning, *Striga* seeds perceive host-derived signals, commonly carotenoid-derived phytohormones called strigolactones (SL), which also shape plant shoot architecture and influence hyphal branching in arbuscular mycorrhizal fungi (Matusova et al., 2005; Gomez-Roldan et al., 2008; Umehara et al., 2008). But germination is in fact a trade-off. On one end, germination in the presence of an appropriate host provides an opportunity to successfully infect a host and complete its parasitic lifecycle. On the other end, germination without an appropriate host leads to certain death after the parasite depletes the limited seed reserves.

Studies of how *Striga* makes this “life or death” decision has expanded our mechanistic understanding of genetic underpinnings of parasite seed germination and in effect provided opportunities for managing the parasite using the approach described as pre-attachment *Striga* resistance. Germination starts with the perception of SLs by a diverse group of α/β hydrolase receptors known as KARRIKIN INSENSITIVE 2/ HYPOSENSITIVE TO LIGHT (HTL/KAI2) (Toh et al., 2015, (Nelson, 2021). Different *Striga* receptors perceive SLs at different rates. In sorghum, for example, *Striga* germinates more effectively in response to 5-deoxy strigol – exuded by most sorghum genotypes relative to a few mutants described as *low germination loci 1* (*lgs1*) that emit orobanchol (Gobena et al., 2017).

Despite these advances, some aspects of *Striga* germination remain unclear and are now the subject of intense investigation. At the forefront is the growing appreciation that the root exudate is a complex blend of biomolecules that could interact to promote or repress *Striga* germination (Mallu et al., 2021, 2022). To shed light on these biomolecular interactions, we used the *S. hermonthica*-sorghum pathosystem in a comparative transcriptome approach that evaluated RNA profiles of high and low inducers of germination against the synthetic strigolactone (R24) positive control. We sought to answer the following questions: (i) what are the genetic factors that underpin *Striga* germination after conditioning? (ii) what are the genetic underpinnings of *Striga* germination in response to high germination inducers relative to low inducers? and (iii) are there synergistic and antagonistic regulators of *Striga* germination in the root exudate? We discuss our results in the context of *Striga* management by exploiting the communication exchange between the sorghum host and the parasite.

(Nelson, 2021)

## 2. METHODS

### 2.1 Plant material

All studies were conducted using *S. hermonthica* seeds obtained from infested fields in Alupe, Kenya (0.45°, 34.13°) in 2018. Sorghum lines i.e., IS2730, IS27146, IS41724, and SRN39 were originally obtained from the International Crop Research Institute for Semi-Arid Tropics (ICRISAT) and are now maintained at the plant transformation laboratory of Kenyatta University, Nairobi, Kenya.

### 2.2 *Striga* germination assay

*Striga* germination was done according to Mwakha et al., (2020), using sorghum root exudate. Sorghum seeds were germinated in a 13.5 cm × 11 cm × 11 cm container set in a complete randomized block design in three replicates. Plants were gently plucked seven days after emergence and each seedling root cleansed and transferred into 50 ml glass tubes with Long Ashton Nutrient medium (Hudson, 1967). Cotton wool plugs were used to support seedlings. Tubes were covered subsequently with aluminium foil to block out light.

Seedlings were left to grow hydroponically in a greenhouse at 28°C during the day (16 hrs) and 24°C at night (8 hrs) under 450 μMm photoperiod and 60 % relative humidity. These seedlings were grown for seven more days and were transferred into clean tubes containing 20 ml sterile distilled water, then wrapped with aluminium foil to be incubated. After 48 hours, roots were weighed, and root exudates collected. Individual root weight was used to normalize the volume of root exudate for each Petri plate.

Root exudate (1 ml) per gram of root weight was used to induce the germination of conditioned *Striga* seeds in each Petri plate. GR24, (Chiralix, Nijmegen, Netherlands) measuring 5 ml at a concentration of 0.1 ppm in double distilled water was used as a positive control while conditioned untreated *Striga* seeds served as a negative control. Petri plates were subsequently sealed using parafilm, covered with aluminium foil, and incubated in the dark at 30°C for 24 hrs.

In the end, six treatments (four root exudates, GR24, and water control) were used to induce *Striga* germination and RNA extracted from each treatment at 6, 12, and 18 hrs after treatment. The experiment had five technical and three biological replicates set in a complete randomized design.

### 2.3 Data analysis

*Striga* seed germination frequency and radical length were measured using ImageJ v. 1.45 (http://rsb.info.nih.gov/ij). The germination frequency was calculated using the equation below:

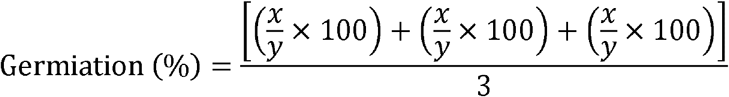

x is the number of germinated *Striga* seeds, y is the total number of *Striga* seeds in the petri dish, and three is the number of replicates in the experiment. Data on seed germination and radical length assessment were determined and the resulting means separated using Turkey’s honest significant difference (HSD) test at a 5 % level of significance. Data was displayed in box plots produced using the ggplot2 package in R (Wickham *et al*., 2019).

### 2.4 RNA extraction

Whole *Striga* seedlings were harvested at three-time intervals (6 hai, 12 hai, and 18 hai). At each time point, seeds were frozen using liquid nitrogen, followed by immediate RNA extraction using a Bioline ISOLATE II RNA Plant Kit (London, UK) according to the manufacturer’s instructions. The obtained RNA samples from each treatment were treated with Deoxyribonuclease I to remove potential DNA contaminations. RNA quantity and quality were determined using a Nanodrop ND-1000 spectrophotometer (Thermo Fisher Scientific), a Qubit fluorometer (Invitrogen), and an Agilent Bioanalyzer. Samples with an RNA integrity number ≥7.0 were subjected to sequencing using the 3’Tag-Seq method as described in Mayer et al. (2011).

### 2.5 Quality control Pre-processing of raw 3′ Tag-RNA-Seq reads

The 3-Tag-RNA-Seq-analysis NextFlow pipeline (https://github.com/fmobegi/3-Tag-RNA-Seq-analysis) which wraps up the TagSeq utilities v2.0 (https://github.com/Eli-Meyer/TagSeq_utilities) was used to process the RNA-Seq data. Briefly, the quality of raw sequences was assessed using FastQC v.0.11.9. Raw reads were filtered based on Phred per-base quality score (Q) ≥ 20 and a cumulative per-read low quality score (LQ) ≤ 10 using the *QualFilterFastq.pl* script with the parameters *-m 20* and *-x 10*. Reads that passed this step were then depleted of homo-polymer repeats longer than 30 bp using *HRFilterFastq.pl* (parameters *-n = 30*), adapter sequences using *Bbduk.sh*, part of *BBMap* v.35.85, and PCR duplicates using *RemovePCRDups.pl*. Non-template sequences introduced at the 5’ end of cDNA tags during Tag-Seq libraries preparation were trimmed using *TagTrimmer.pl* (parameters: *-b 1 -e 8*)

Quality-processed reads were re-evaluated for quality using FastQC and then mapped using hisat2 (Kim *et al*., 2019) to the *S. hermonthica* transcriptome reference accession numbers SRX040928, SRX040929, SRX008132, and SRX008133. SAMtools v.1.10 was used to convert the generated sequence alignment map (SAM) files into binary alignment mapping (BAM) format, and to sort and index the BAM files. The sorted BAM files were used as input for transcripts quantification using Salmon (Patro *et al*., 2017), a quasi-mapping-based and quantification method, against the *S. hermonthica* transcript reference. The ensuing raw counts in form of TPM were collated in R using the Bioconductor package tximport (Soneson et al., 2016) (parameters; *countsFromAbundance* = “*lengthScaledTPM*”) and exported for downstream differential gene expression analysis.

### 2.6 Analysis of differential gene expression

Raw counts were transformed into counts per million (CPM) using the *cpm()* function in *edgeR* package (Robinson et al., 2009). Genes that had a CPM of greater than 1 were retained for further analysis. Principle coordinate analysis (PcoA) plots and heatmaps were generated using *ggplot2* to determine the relatedness of the biological replicates. Differential gene expression analysis was performed using *DESeq2* Bioconductor R package [(Love *et al*., 2014) (http://www.bioconductor.org/packages/release/bioc/html/DESeq2.html] following normalization using *DESeq2* with a Wald test. A false discovery rate (FDR) cut-off of 0.05 was applied, and a log_2_ fold change cutoff of ≥2 to indicate upregulation and ≤−2 to indicate down-regulation. Differentially expressed genes (DEGs) were considered at each time point for each host in relation to the controls.

Gene IDs were used to obtain fasta files of interest from the *S. hermonthica* transcriptome using customized Bio Python scripts and then annotated using *Blast2GO* (Conesa et al., 2005). Resulting Gene ontology (GO) terms were derived from InterPro and the Kyoto Encyclopaedia of Genes and Genomes (KEGG) were then subjected to pathway enrichment using ShinyGO (Ge *et al*., 2019). Pathway enrichment analysis of DEGs were performed using the R Bioconductor package *TopGO* implementing the classical method and Fisher’s exact test with a *P*-value threshold of ≤0.05 (reference: https://academic.oup.com/bioinformatics/article/22/13/1600/193669). Differential gene expression was visualised using volcan plots and upset plots produced using ggplot2 (Wickham, 2016) and PlotUpset (Conway et al., 2017) in R. Differentially expressed genes hierarchical clustering was performed using 4×2 hexagonal SOMs using the package kohonen in R (Wehrens and Kruisselbrink, 2018).

## 3. RESULTS

### 3.1 *Striga*’s variable germination response to different sorghum genotypes

Germination rate of *Striga* is variable depending on various blends of SL emitted by hosts (Gobena et al., 2017). Therefore, we first measured the germination efficiency and radical lengths of the selected sorghum genotypes: SRN39, IS41724, IS27146 and IS2730 evaluated against a positive control (synthetic SL, GR24).

Germination efficiencies and radical lengths observations were made at 6, 12, and 18 hours after induction (hai) as described in Figure 1a. Analysis of germination efficiencies and radical lengths are shown in Figures 1b and c. We found that germination had occurred at 6 hai in IS2730 (7.18%), GR24 (2.32%), and IS27146 (1.99%) but not in SRN39 and IS41724. At 12 hai, germination had occurred in all genotypes, and it was higher in GR24 (28.14%) than in IS2730 (24.02 %), IS27146 (15.44%), IS41724 (15.07%), and SRN 39 (21.60). At 18 hai, germination increased in all genotypes. Overall, there were no differences in seeds treated with IS2730 (32.16%) and GR24 (29.59%), but these values were higher than IS27146 (23.78%), SRN39 (24.61%), and IS41724 (20.15%) observed at that time point. Radical length differences also varied across all treatments at different timepoints. At 6 hai, radical growth was observed in GR24 (0.058mm), IS2730 (0.092mm), and IS27146 (0.089mm). At 12 hai, Turkey’s mean separation using HSD resulted in two groups of radical lengths that were significantly different. GR24 (0.19mm) and IS41724 (0.18mm) had longer radical lengths than IS2730 (0.15mm), IS27146 (0.12mm), and SRN39 (0.13mm). At 18 hai, *Striga* seeds in all treatments had similar radical lengths except seeds treated with IS2730 which had longer radicals (0.56 mm). These results underscore significant variations in germination induction kinetics by the genotypes at the different time points.

**FIGURE 1.**
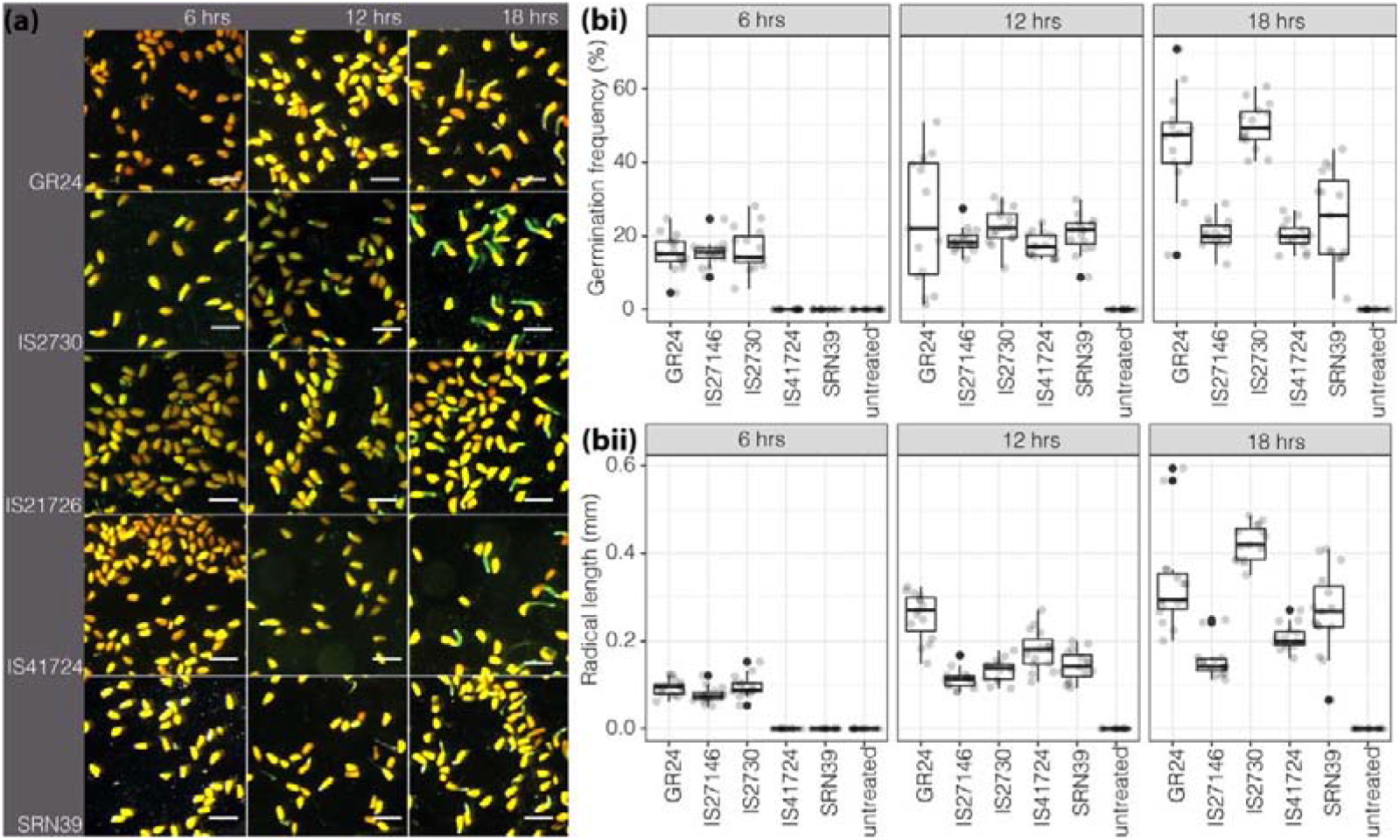
Germination response of *Striga* to different root exudate treatments. (a) Illustrative images of *Striga* germination induction using treatments of root exudates from different sorghum genotypes compared with the synthetic strigolactone (GR24) and SRN39 as positive and susceptible checks. Induction was performed at 6, 12, and 18 hours of induction. Scale bar = 1 mm. (b) Rates of *Striga* germination as induced by various treatments at 6, 12, and18 hours after induction (hai). At 6hai, germination was already observed in high germination inducers and the positive control but not in low germination inducers (IS41724 and SRN39). *Striga* radical lengths after treatments using root exudates of various sorghum genotypes at different time points. Early induction of germination by high inducers of germination led to notably longer radical lengths.

### 3.2 Transcriptional gene activation underlies rates of *Striga* germination induction by sorghum genotypes

To examine the genetic complexity of observed germination rates, we performed a comparative transcriptome analysis of germination induction of the sorghum genotypes at the 6 hai, 12 hai, and 18 hai.

Comparing the number of DEGs with germination rates of the different treatments pointed to a correlation between germination induction and the number of DEGs, implying that an earlier activation could be responsible for the different germination rates in the treatments (Figure 2 and Figure S1). In support of this hypothesis, there was a notable increase in DEGs for SRN39 and IS41724 at 12hai (when these genotypes induced germination), and a decrease in DEGs in seeds induced by IS2730 and GR24 treatments at 12 hai except for the number of DEGs in IS27146. At 18 hai, DEGs had decreased in all treatments except in SRN39. Figure 2 further show common genes whose expression was in all treatments at 6hai, at 12hai, and at 18 hai. GO enrichment for the intersect of genes at 6 hai showed that these genes are involved in cell wall loosening, amino acid metabolism, carbohydrate metabolism, and energy metabolism. At 12 hai, DEGs were enriched for cell wall loosening and catabolism of cell wall material (arabinian and xylan metabolism) while at 18 hai, they were enriched for (polyribonucleotide nucleotidyltransferase) HSP, glucose-repressible protein, oil body-associated protein 2C-like probably involved in seed growth. We presumed that the genes that were differentially expressed in all treatments were part of core germination processes.

**FIGURE 2.**
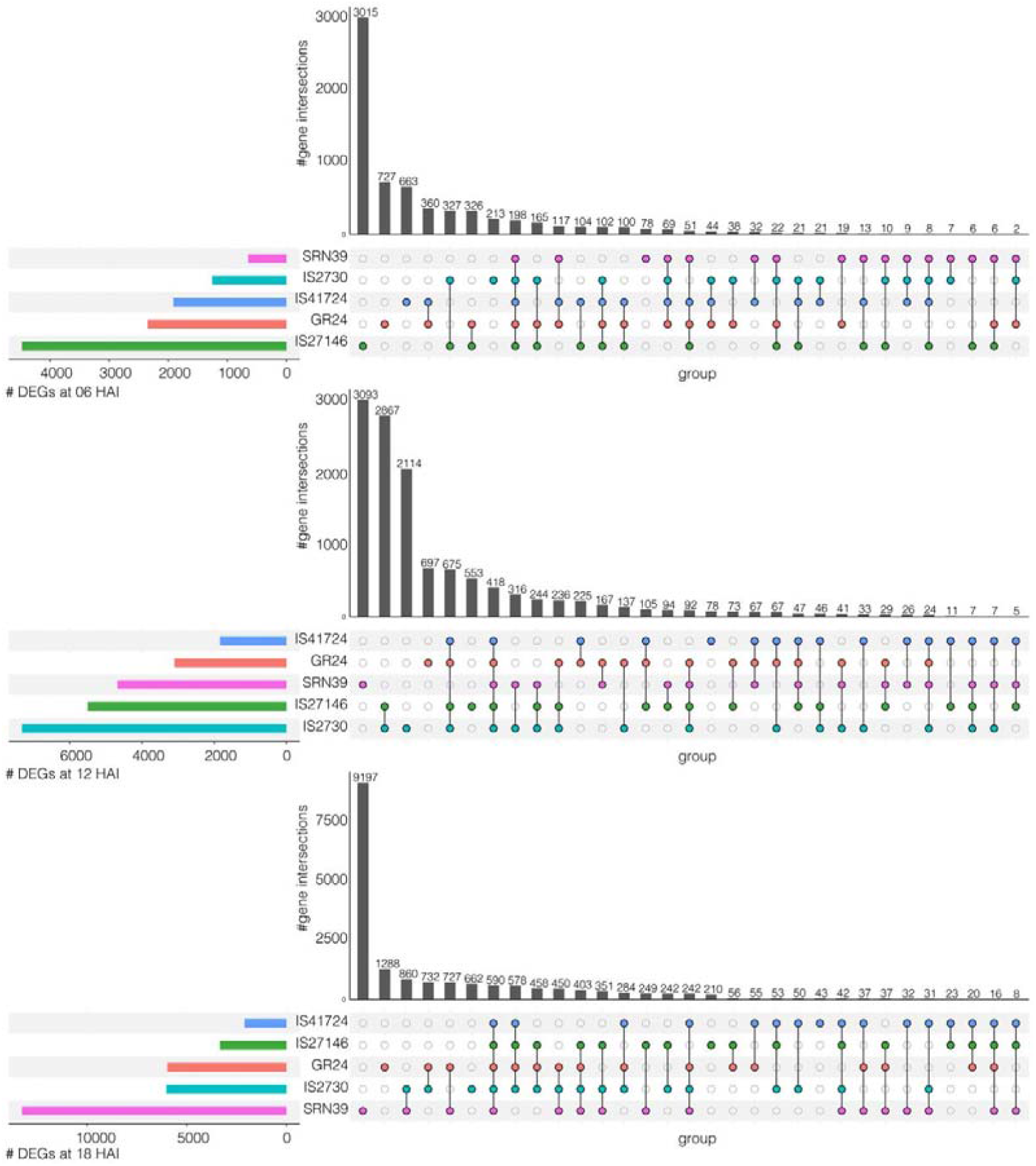
Transcriptional reprogramming during *Striga* germination. An upset plot showing the overlap of differentially expressed genes (DEGs) between comparisons. The coloured horizontal bar graphs indicate the number of DEGs for each comparison, respectively. The black vertical bar graphs show the intersection size of DEGs and the colored dots represent contributing comparisons from the DE analysis. For fast inducers of germination, many genes are induced as early as 6 hrs (upper panel on IS27146). Gene induction is also associated with gene induction at 12 hai (IS2730 middle panel) and at 18 hai (SRN39, lower panel) for late inducers of germination. Gene induction for germination stimulation is specific to genotypes with some unique genes only getting induced by specific germination stimulants, but there is also conservation where common core genes are induced by different treatments.

### 3.3 *Striga* germination is phased, and genotype differentiated

To determine the dynamics of germination induction responsible for variability in *Striga* germination rates as induced by various sorghum genotypes, we categorized the treatments by hierarchical clustering using self-organizing maps (SOMs). Principal component analysis (PCA) and line graphs for scaled gene expression chats are shown in Figure 3a and 3b. There were 8 clusters that differed in the gene expression patterns. Notably, the expression pattern in SOM5 showed a steady increase in gene expression with a clear differentiation between imbibed non-germinated seeds and germination induction treatments at 6 hai. In these clusters, gene expression went up at 12 hai but declined at 18 hai. Other notable SOM clusters were SOM4, SOM7, and SOM8 which showed either genotype-specific or a time-specific gene expression pattern. Enrichment pathways of these SOM clusters showed specific developmental processes of seed germination. This is well illustrated with SOM8 in relation to SOM4. SOM8 represents the pathways that defined a switch between dormancy break and germination. This SOM was significantly enriched for response to Karrikin – representing response to the germination stimulants, followed by notable DNA repair processes, dormancy release process, and embryo development. Consistent with the progression of the germination process, SOM4 was enriched for protein synthesis using extant mRNAs, seed reserve mobilization, further DNA repair, transport of solutes, and notably, seedling development. The low activity in 12 and 18 hai in the early germinators is consistent with repression of seed maturation to allow a transition to germination (Bewley, 1997). A similar inference can be made regarding genotype specificity in germination. Early induction of gene expression was in GR24, IS2730, and IS21746 as indicated in SOM4 induction of seed maturation pathways. Induction of genes for IS41724 and SRN39 was highest at 12 hai and 18 hai. Pathways induced in this cluster included an overlap between early induction of germination processes and later germination processes. Similarly, SOM7 was specific to induction of germination by SRN39 and IS41724 – indicating similarities in these two genotypes. Because both SRN39 and IS41724 are low inducers of germination, we presumed that the similarities could be associated with the resistant phenotype. Together, these results point to a time-differentiated response to germination stimulation by the different treatments. The rates are likely dependent on the perception of germination signals in the different treatments.

**Figure 3.**
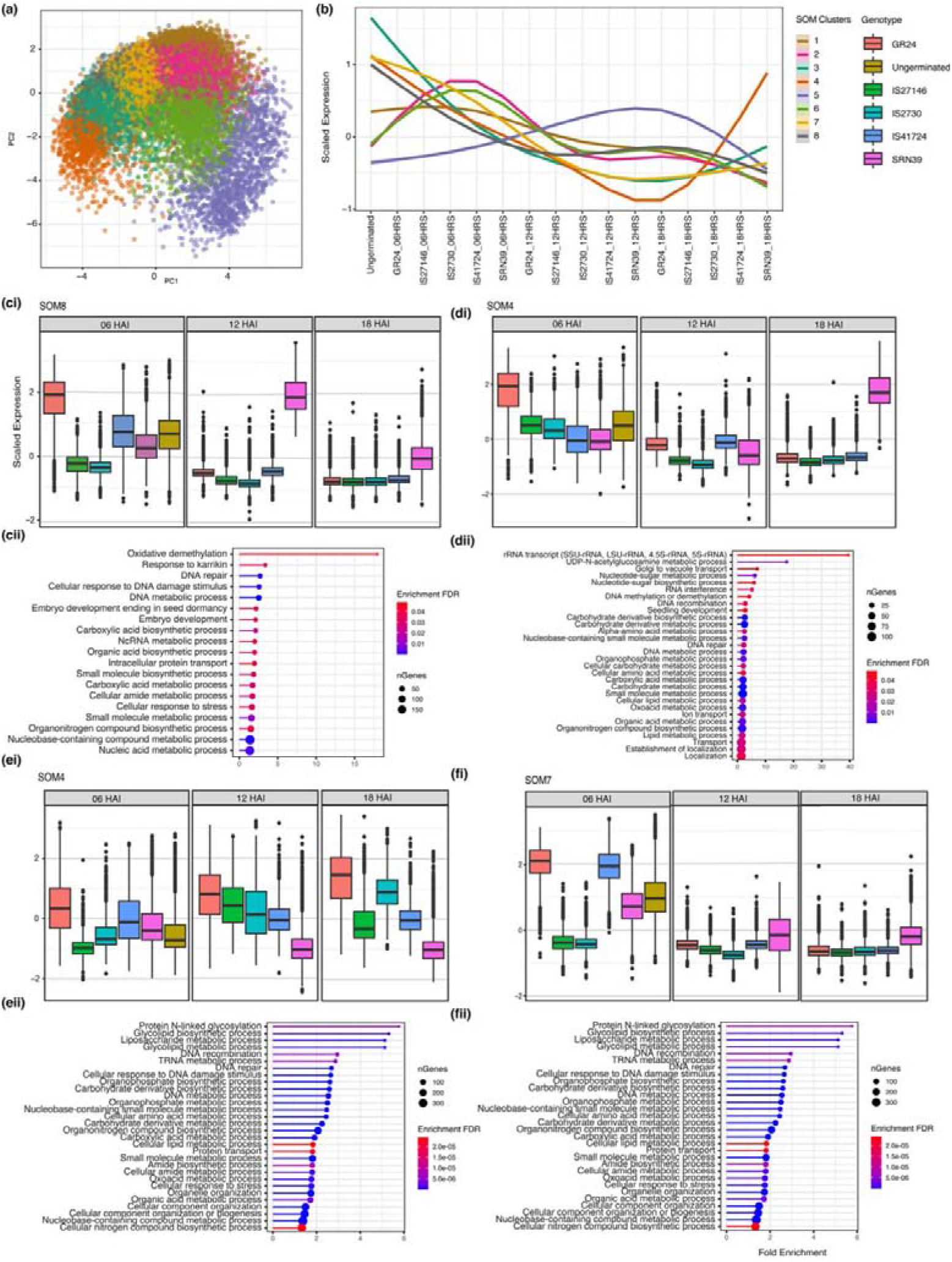
Time and sorghum genotype differentiated clusters of *Striga* germination induction. (a) PCA of a 4×2 SOM for various genotypes and times of induction and root exudate vs. GR24 showing 8 clusters. (b) Line graphs showing trends of gene induction and repression and relationships between SOMs. SOMs 4, 5, 7, and 8 showed contrasting gene expression patterns for the genotypes and the times after induction. (ci) gene expression patterns for SOM8 and GO enrichment (cii) showing a phased gene expression for early germination processes compared to late gene expression patterns (di) and pathways (dii). Gene expression patterns in SOM 4 for showing similarity in all treatments at early germination (ei) and accompanying enrichment analysis compared to genotype-specific expression of SRN39 and IS41724 (fi) and enrichment in (fii).

### 3.4 *Striga* germination is associated with Abscisic acid repression

The switch from dormancy to germination occurred at 6 hours upon perception of the germination signal – SL. Studies of seed germination have implicated the hormone abscisic acid (ABA) as the primary growth regulator for dormancy release (Gianinetti and Vernieri, 2007; Nambara *et al*., 2010; Shu *et al*., 2016). We therefore explored the hypothesis that germination occurred with repression of ABA against the alternate that ABA has no role in influencing *Striga* germination rates. We analysed specific gene expression patterns in the ABA biosynthesis pathway mapped in the carotenoid biosynthesis pathway of the Kyoto Encyclopaedia of Genes and Genomes (KEGG). The results are shown in Figure 4.

**FIGURE 4.**
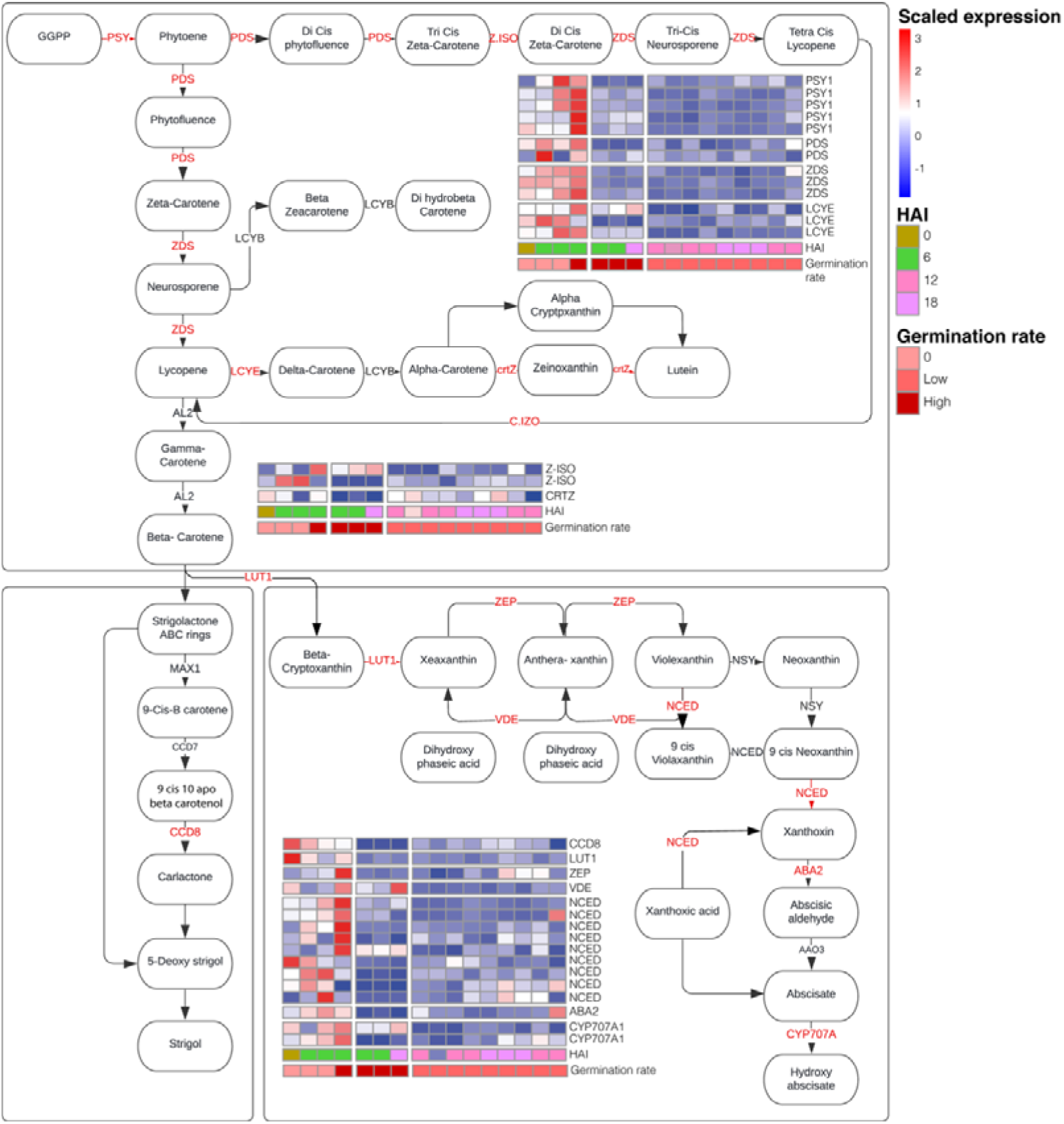
Expression of carotenoid biosynthesis genes. ABA biosynthesis genes were induced at 6 hai and repressed at 12 and 18 hai. Repression of ABA corresponded with germination.

The first step in carotenoid biosynthesis is the conversion of phytoene lycopene. The enzymes involved in this step PSY, PDS, Z-ISO, ZDS, and CRTISO were all consistently induced in non-germinated seeds, and treatments of resistant sorghum and GR24 treated seeds at 6 hai. Lycopene is then catalysed into carotene by the cyclases, LCY-B and LCY-E. In our case, LCY-E was significantly induced in the non-germinating seeds, resistant sorghum and GR24. Finally, carotenoids are degraded by carotenoid cleavage dioxygenases (CCD) to form either SLs or ABA. Three CCD genes, and four NCED genes were all induced in non-germinating seeds, resistant treatments and GR24 at 6 hai. In all cases, germinating seeds showed repression of the carotenoid biosynthesis and degradation genes. Because of the role of the enzymes encoded by these proteins in the biosynthesis and degradation of carotenoids, we conclude that the observed changes in gene expression may cause differences in germination rates. Furthermore, because of inhibitory roles of ABA in seed germination, we surmised that ABA repressive effects must be alleviated for seed germination to occur. But appearing to contradict this hypothesis is notable induction of carotenoid biosynthesis genes in germinating seeds treated with GR24. This could be suggestive of ABA suppression by components in root exudate but and not in GR24. Equally perplexing is the induction of the SL biosynthesis gene CCD8 that leads to biosynthesis of the intermediate carlactone. Bearing in mind that *Striga* lacks the downstream genes that lead to canonical strigolactone biosynthesis, it was not clear why this enzyme was induced (Abe et al., 2014). One possibility is that CCD8 expression may just be responding to increased traffic down the SL biosynthesis path when carotenoid biosynthesis is high. The observations however bring to the fore a possible interaction between SL, ABA and other hormone pathways.

### 3.5 *Striga* germination is modulated by a hormonal crosstalk

To address the question of a plausible hormonal crosstalk modulating *Striga* germination, we mapped the differentially expressed genes during *Striga* germination as induced by different treatments on the hormone signaling pathways in KEGG maps. We considered genes in the ABA, auxin, brassinosteroid (BS), cytokinin (CK), ethylene (ET), gibberellic acid (GA), jasmonic acid (JA) pathways (Figure 5). Analysis of gene expression patterns showed: ABA and auxin repressed germination. ABA genes PPC2, SnKR2, and ABF were repressed in germinating seeds and induced in non-germinating seeds. This was also the case for auxin genes (AUX1, TIR1, GH3, ARF and SAUR). The observed gene expression patterns are consistent with mechanistic of germination inhibition by ABA and auxin. ABA germination suppression occurs upon activation of PP2C by ABA, triggering release of SnRK2 kinase, and subsequent activation of ABSCISIC ACID INSENSITIVE5 (ABI5) that exerts transcriptional control over seed dormancy (Fujii et al., 2009). Likewise, auxin represses germination through ARF through stabilization of the ABA transcription factor ABI3 (Liu et al., 2013).

**FIGURE 5.**
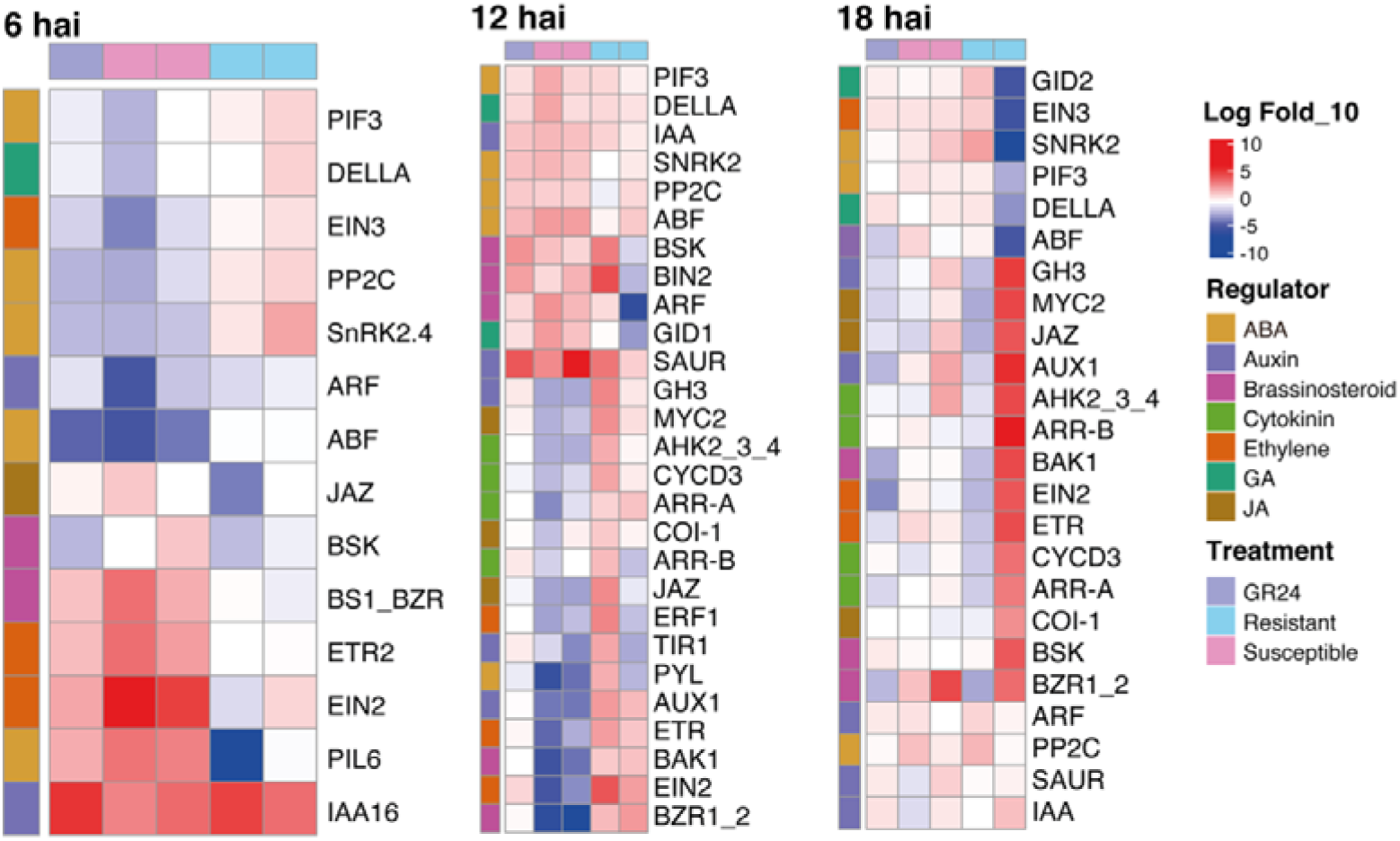
The role of hormones in the Striga germination temporal patterns. Heatmaps show gene expression patterns of the plant hormone signaling pathways of Kyoto Encyclopaedia of Genes and Genomes (KEGG) map at 6, 12, and 18 hai as induced by different root exudate and GR24 treatments.

Brassinosteroid, ET, GA and JA enhanced germination as follows: (i) BS genes BS1, BSK BZR and BAK1 were induced in germinating seeds pointing to a positive regulatory of seed germination through BSs. It is known that BIN2 interact with ABI5 to mediate antagonism between ABA and BS (Hu and Yu, 2014) and that SL-dependent *Striga* germination requires functional BS signaling (Bunsick and Lumba, 2020). (ii) Ethylene proteins (ETR, ERF and EIN2) were consistently induced in germinating seeds reinforcing the extensively studied antagonism of ethylene on ABA reviewed in (Linkies and Leubner-Metzger, 2012; Gazzarrini and Tsai, 2015). Furthermore, PIL6 which promotes ethylene activity in the dark (Khanna et al., 2007) was induced in germinating seeds in a pattern like ethylene proteins. (iii) GA repressor gene DELLA was repressed in germinating seeds at 6 hai. Although this pattern points to a positive regulatory role of GA in *Striga* germination, DELLA was induced in germinating seeds at 12 hai, making it difficult to specifically determine if GA indeed has a positive regulatory role in *Striga* seed germination. Finally, (iv) JA, signaling genes JAZ, MYC and CO1 were induced in germinating seeds indicating that JAZ is a positive regulator of germination in agreement with previous works demonstrating JAZ’s repression of ABI3 and ABI5 (Pan et al., 2020).

We considered that CK has a less prominent role in *Striga* germination on account that there were no cytokine DEGs at early germination stages. Cytokinin genes A-ARR and B-ARR were upregulated in already germinated seeds at 12 and 18 hai leading us to speculate that these genes had functions after germination. Because of notable expression of PIF3 transcription factors involved in photomorphogenesis (Kim et al., 2003; Feng et al., 2008), and cell cycle genes (CYDC) we deduced that CK gene activation was involved phytochromes interactions of ketomorphogenesis. Interestingly, CK specifically antagonizes ABA-mediated inhibition of cotyledon greening with minimal effects on seed germination in (Guan et al., 2014) further affirming a possible post-germination, rather than a germination role for CK.

In summary, our results suggest that: (i) ABA and Auxin repress germination, (ii) BS, ET and JA promote germination; (iii) CK and GA may have a more prominent role post-germination rather than during germination. We provide a proposed model in Figure 6.

**FIGURE 6.**
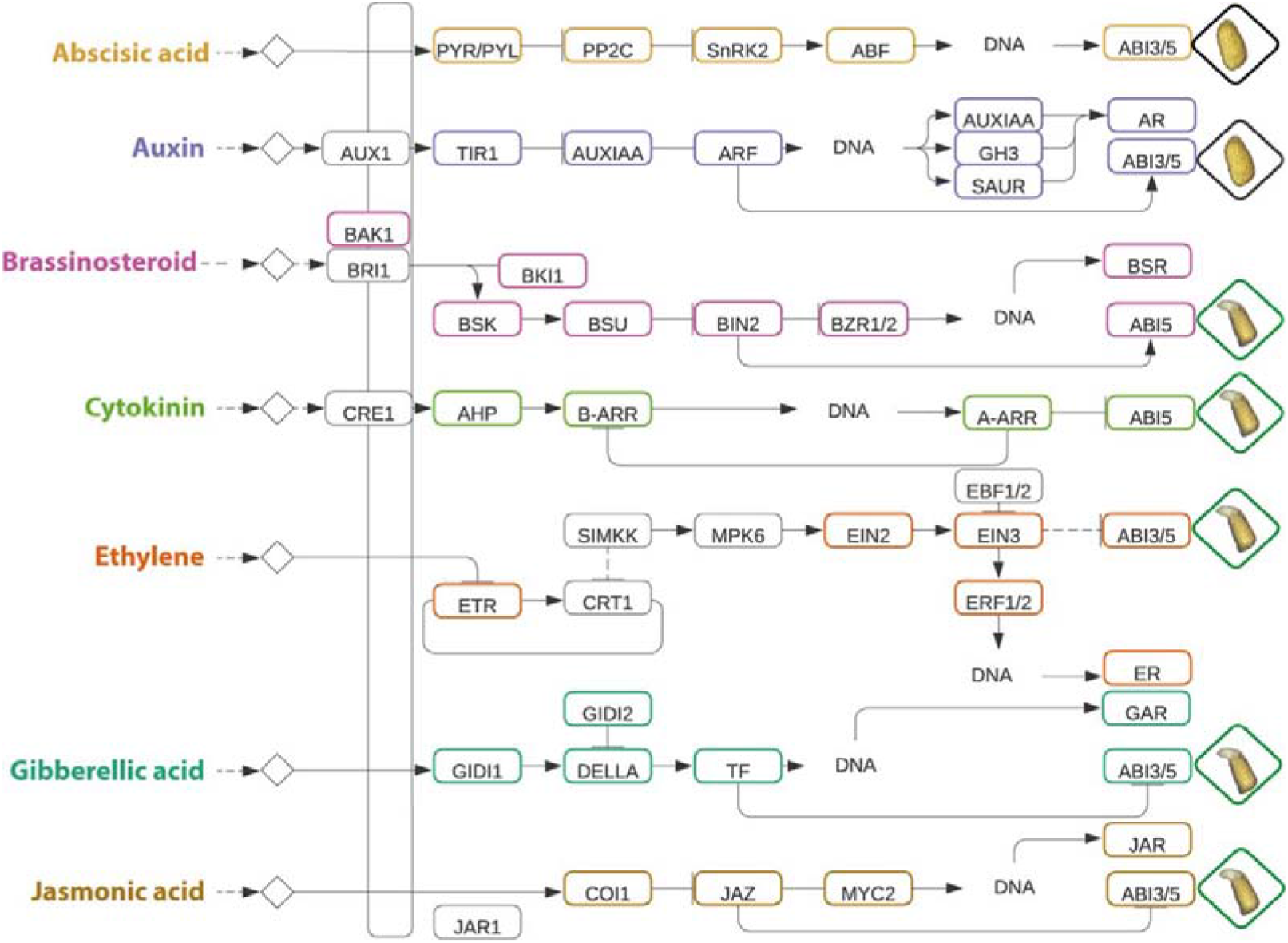
Proposed model of hormone signaling moderation based on gene expression patterns of the Kyoto Encyclopaedia of Genes and Genomes (KEGG) map for plant hormone signaling highlighting positive and negative regulators of *Striga* seed germination. Differentially expressed genes for each pathway are color-coded.

## 4 DISCUSSION

Processes that lead to germination of the parasitic plant *Striga* could provide useful insights into strategies for its management. Principally, *Striga* germinates in response to germination stimulants emitted from the host root exudate. However, germination is highly variable in most *Striga*-host interactions. In sorghum for example, genotypes that have a mutation on the *lgs1* loci – SRN39-like emit the less powerful germination inducer SL (orobanchol) unlike their *LGS1* counterparts that emit the powerful inducer 5-deosystrigol. Genotypes (*lgs1*-like) stimulate *Striga* seed germination less efficiently and are advocated for *Striga* resistance breeding in SSA. To unravel the genetic basis of this complexity in sorghum, we did a comparative transcriptomic analysis of the various genotypes of sorghum that exhibit variation in stimulating *Striga* germination. We sought to identify not only the major transcriptomic changes that define the germination process in *Striga* but also a set of genes that may contribute to the differences in germination rates exhibited by these genotypes. For this purpose, we carried out RNA sequencing (RNA-seq) on four selected genotypes (IS2730, IS27146, IS41724, and SRN39) and the positive control (synthetic SL, GR24) at different stages of seed germination.

Analysis of differential gene expression of *Striga* seeds in response to stimulation by different root exudates revealed distinct genotypic differences and core genes required for germination. The core DEGs were enriched for amino acid metabolism, carbohydrate metabolism, energy metabolism, biosynthesis of other secondary metabolites, lipid metabolism, and signal transduction. This is consistent with the metabolic processes of seed germination in other studies (Weitbrecht et al., 2011; Boter et al., 2019). Studies have shown that the seed expands rapidly at the initial stage of germination due to water absorption. Genes associated with sugar, lipid and amino acid metabolism are upregulated. Additionally, cell wall synthesis and degradation genes are upregulated to provide small molecules and materials for cell wall synthesis during seed germination (Boter et al., 2019).

We then used self-organizing maps to cluster differentially expressed genes according to the rates of germination induction by various treatments at different time points. The switch to germination occurred first at 6 hai. This timepoint marked the dormancy release phase and commencement of germination. Germination itself was phased into early and late germination processes. Early germination was characterized by the perception of SL, resource mobilization, DNA repair, dormancy release, and embryo development while late germination was characterized by protein synthesis and carbon fixation. Germination was also genotype-specific, resulting in early and late inducers of germination. Rapid seed maturation processes were observed in the GR24, IS27146, and IS2730 treatments. Late seed maturation was observed in seeds treated with root exudates of SRN39 and IS41724, and IS27146. SOM analysis showed that early germination (6hai) processes in early inducers were clustered with later treatments of later inducers, implying that these processes were delayed in late inducers. This finding is consistent with what is known about germination induction by IS41724 and SRN39. Previous work has shown that these genotypes support poor *Striga* germination and radical growth (Mallu et al., 2021, 2022). Our findings are also consistent with previous studies of seed germination. Transcriptomic analysis of barley seeds (Leišová-Svobodová et al., 2020) revealed transient upregulation of cell wall synthesis genes. Regulatory components such as transcription factors, signaling, and post-transcriptional components are transiently upregulated during the early germination phase, whereas the late germination phase is characterized by transcriptional activation of genes representing the histone families and many metabolic pathways such as amino acid metabolism, nucleotide metabolism, and those related to cell division and cell cycle. The post-germination phase is characterized by the induction of genes involved in photosynthesis and reserve mobilization.

Because the carotenoid pathway leads to the biosynthesis of *Striga* germination crucial phytohormones, SL and ABA, we examined differential gene expression patterns in the different treatments for a possible modulation of germination because of these hormones. Consistently, ABA was induced in non-germinating, resistant, and GR24 treatments in the early phase of germination induction (6 hai) but repressed in germinating seeds and late stages of germination induction (12 and 18 hai). This observation is reminiscent of an ABA dormancy release phenomena previously observed in studies of seed germination (Kusumoto et al., 2006; Mallu et al., 2022). Our study also pointed to a possible interaction between ABA and other hormones.

Therefore, the last aspect of our study investigated the role of phytohormones in regulating the rates of *Striga* germination. Induction of ABA and auxin genes corresponded with the suppression of germination. This is consistent with physiological germination experiments that show ABA severely limits *Striga* germination, and its repeal using the inhibitor fluridon causes spontaneous germination of *Striga* seeds (Kusumoto et al., 2006; Mallu et al., 2022). Although not shown in *Striga*, the role of auxin in dormancy imposition in an ABI3-dependent manner has been well described in Arabidopsis (Liu et al., 2013). This is also consistent with literature that show that auxin leads to a release of ARF, which stabilizes the binding of ABI5 and thereby enhances ABA responses. Both processes lead to the inhibition of germination in an ABI5-dependent manner. Enhanced germination in interactions with other hormones such as BS and JAZ could result from antagonizing ABI5. Such a hypothesis is supported by the fact that ABI5 physically interacts with the BRASSINOSTEROID INSENSITIVE2 (BIN2) and brassinosteroid-related protein kinases, and that the BIN2-ABI5 cascade mediates the antagonism between BS and ABA during seed germination (Hu and Yu, 2014). Additionally, JAZ proteins directly interact with ABI5 to negatively regulate ABA (Ju et al., 2019). Ethylene is known to play a role in dormancy release (Corbineau et al., 2014) and spontaneous germination of *Striga* seed (Logan and Stewart, 1991). Regarding GA, there are reports on GA being a positive regulator of *Striga* germination (Seo et al., 2008) while others have reported GA as a negative regulator of *Striga* germination (Ito et al., 2017; Mallu et al., 2022). We found that DELLA, a GA repressor, was downregulated in germinating seeds, but upregulated in non-germinating seeds. Unlike seeds of other plants, bypasses GA-dependent germination in favour of SL-dependent germination (Bunsick et al., 2020) and this may confound results of exogenous GA application.

Curiously, we observed a notable association of hormones with the phytochrome-interacting transcription factor (PF3) post-germination. Under light conditions, PF3 negatively regulates light responses, including hypocotyl elongation, cotyledon opening, and hypocotyl-negative gravitropism. This brings to focus the remarkable capability of *Striga* and other parasitic plants to suppress shade phenotypes – elongated hypocotyl growth and closed cotyledons. One possibility is that *Striga* maintains hyperactive phytochromes consistent with the phenotypes of PHYA over-expressors (FRANKLIN and WHITELAM, 2005). Interestingly, it was recently shown that ABA modulates shade avoidance by inducing hyponasty movement in Arabidopsis (Michaud et al., 2022) adding to another piece of circumstantial evidence for ABA involvement in *Striga*’s germination and infection process. This is an interesting subject for future investigation.

In conclusion, we have shown that *Striga* seed germination is controlled by a complex signaling and a gene expression regulatory network. Perception of the germination signal (SL) sets the germination program for different rates of germination or abortion. After SL perception, *Striga* appears to share similar molecular mechanisms, such as phytohormone behaviour, transcription and translation activation, and the process of radical protrusion, with other plants. Beyond germination, hormones interact with phytochromes to ensure *Striga* seedling growth and development sub-terranean.

## ACKNOWLEDGEMENTS

This work was supported by the National Academies of Science (NAS) under the Partnerships for Enhanced Engagement in Research (PEER) program (contract number NAS Sub-Grant Award Letter Agreement Number 2000011208). The Sub-Grant Agreement was funded under Prime Agreement Number AID-OAA-A-11-00012, which was entered into by and between the NAS and the United States Agency for International Development (USAID). Transcriptomic sequencing was supported by funds from the Arkansas Biosciences Institute (the major research component of the Arkansas Tobacco Settlement Proceeds Act of 2000). We further thank the East African Community Scholarship program funded by the German Development bank and implemented by the Inter-university Council of East Africa, and Adroit international for supporting Irafasha Gilles Masters fellowship. Finally, we thank the Alexander von Humboldt Foundation’s George Forster Fellowship for Seniors researchers for supporting Steven Runo’s sabbatical leave.

## AUTHOR CONTRIBUTIONS

S.R., E.S.B. designed the study; I.G., S.M., B.H., collected data; F.M., S.R. and I.G. analysed the data with help from E.S.B, A.W. and S.W.; I.R. wrote the manuscript with help from F.B., S.R., I.G. and S.R., S.W., G.O., and E.S.B.

## DATA AVAILABILITY STATEMENT

Raw reads are available from the National Center for Biotechnology Information Sequence Read Archive under Bioproject accession number PRJNA904327.

---

**Fig. S1.**
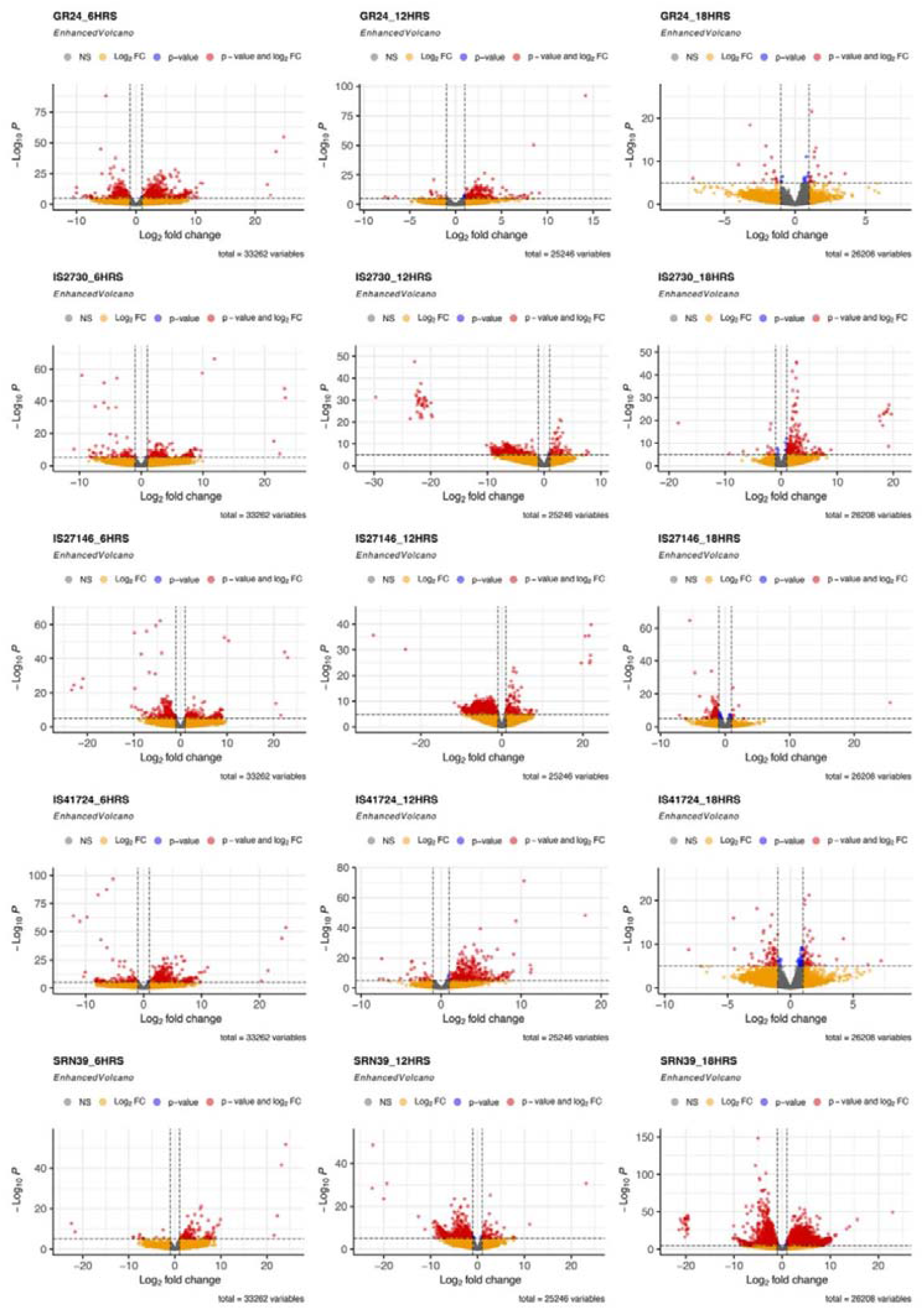
Transcriptional gene activation underlies rates of *Striga* germination induction by sorghum genotypes. Volcano plots representing numbers of differentially expressed genes in germinated *Striga* seeds as induced by root exudates of various sorghum genotypes at 6, 12 and 18 hours. Transcriptional reprogramming marked the start of germination. This occurred at 6 hai for early inducers but 12 hours for later inducers of germination.

